# HIV-1 Vpu promotes phagocytosis of infected CD4^+^ T cells by macrophages through downregulation of CD47

**DOI:** 10.1101/2021.03.16.435750

**Authors:** Lijun Cong, Scott M. Sugden, Pascal Leclair, Chinten James Lim, Tram NQ. Pham, Éric A. Cohen

## Abstract

Human immunodeficiency virus (HIV) remodels the cell surface of infected cells to facilitate viral dissemination and promote immune evasion. The membrane-associated Vpu accessory protein encoded by HIV-1 plays a key role in this process by altering cell surface levels of multiple host proteins. Using an unbiased quantitative plasma membrane profiling approach, we previously identified CD47 as a putative host target downregulated by Vpu. CD47 is a ubiquitously-expressed cell surface protein that interacts with the myeloid cell inhibitory receptor SIRPα to deliver a “don’t-eat-me” signal, thus protecting cells from phagocytosis. In this study, we investigate whether CD47 modulation by HIV-1 Vpu might promote the susceptibility of macrophages to viral infection via phagocytosis of infected CD4^+^ T cells. Indeed, we find that Vpu downregulates CD47 expression on infected CD4^+^ T cells leading to an enhanced capture and phagocytosis by macrophages. Interestingly, it is through this process that a CCR5-tropic transmitted/founder (T/F) virus, which otherwise poorly infects macrophages in its cell-free form, becomes infectious in macrophages. Importantly, we show that HIV-1-infected cells expressing a Vpu-resistant CD47 mutant are less prone to infect macrophages through phagocytosis. Mechanistically, Vpu forms a physical complex with CD47 through its transmembrane domain and targets the latter for lysosomal degradation. These results reveal a novel role of Vpu in modulating macrophage infection, which has important implications for HIV-1 transmission in early stages of infection and the establishment of viral reservoir.

**IMPORTANCE:** Macrophages play critical roles in HIV transmission, viral spread early in infection, and as a reservoir of virus. Selective capture and engulfment of HIV-1 infected T cells was shown to drive efficient macrophage infection suggesting that this mechanism represents an important mode of infection notably for weakly macrophage-tropic T/F viruses. In this study, we provide insight into the signals that regulate this process. We show that the HIV-1 accessory protein Vpu downregulates cell surface levels of CD47, a host protein that interacts with the inhibitory receptor SIRPα to deliver a “don’t-eat-me” signal to macrophages. This allows for enhanced capture and phagocytosis of infected T cells by macrophages, ultimately leading to their productive infection even with T/F virus. These findings provide new insights into the mechanisms governing the intercellular transmission of HIV-1 to macrophages with implications for the establishment of the macrophage reservoir and early HIV-1 dissemination *in vivo*.

## INTRODUCTION

The viral protein U (Vpu) is a membrane-associated accessory protein encoded by human immunodeficiency virus type 1 (HIV-1) and related simian immunodeficiency viruses (SIVs) but not by HIV-2. Consistent with the roles of HIV-1 accessory proteins in targeting cellular restriction factors to favor immune evasion and viral dissemination, Vpu counteracts many host proteins including BST2/Tetherin to promote efficient viral particle release (1, 2) and CD4 to avoid superinfection and subsequent premature cell death (3). The downregulation of both CD4 and BST2 also protects HIV-1 infected CD4^+^ T cells from antibody-mediated cellular cytotoxicity (ADCC) (4). Given the contribution of Vpu towards HIV pathogenesis, partly through targeting BST2 and CD4, there is a continuing interest in identifying additional Vpu targets. To date, a diverse list of host factors has been identified including CD1d, NTB-A/SLAM6, PVR/CD155, CCR7, CD62L; SNAT1 (5), ICAM1/3 (6), CD99 and PLP2 (7), PSGL-1 (8), Tim-3 (9), and there likely are more to be discovered. Indeed, using a stable isotope labelling of amino acids in cell culture (SILAC)-based proteomic approach, we and others previously identified CD47 as a potential target that is downmodulated by Vpu (6, 10).

CD47, also known as integrin-associated protein (IAP), is a ubiquitously-expressed type I transmembrane protein (11) that serves as a ligand of the signal regulatory protein-alpha (SIRPα, or CD172a), an inhibitory receptor mainly expressed on myeloid cells, like macrophages and dendritic cells (DCs) (12, 13), but also on cytolytic T lymphocytes (14). The interaction between these two proteins results in a “don’t-eat-me” signal that inhibits phagocytosis of target cells expressing CD47 by macrophages and DCs, thus providing an important regulatory switch for the phagocytic function of these cells.

Macrophages make up a heterogenous population of immune cells that play important roles in tissue homeostasis and host defense against pathogens partly through their phagocytic function (15). They are increasingly recognized as important cellular targets of HIV-1 infection (16, 17). Indeed, given their relative lengthy life span and unique ability to resist HIV-1 cytopathic effects and CD8^+^ T cell-mediated killing (18, 19), macrophages are thought to be an important viral sanctuary and vector for HIV-1 dissemination as well as a potential viral reservoir during antiretroviral therapy (ART) (20–23). Macrophages are among the early targets of HIV-1 infection given their proximity to the portal of viral entry, commonly the mucosal tissue (21, 24). They express the CD4 receptor and both chemokine coreceptors CXCR4 and CCR5. While macrophage-tropic (M-tropic) viruses mainly use CCR5 as coreceptor, they are paradoxically mostly isolated from brain tissues of AIDS patients at late stages of infection (17, 25). Yet Infected tissue macrophages can be detected at all stages of disease (26) and T/F viruses that initiate infection as well as inter-individual transmission are weakly M-tropic (27). It is therefore crucial to understand the mechanisms by which macrophages become infected during the early phase of infection. In this regard, it has been reported that macrophages capture SIV- or HIV-1-infected T cells, retain infectious particles in a non-degradative compartment and ultimately become infected (28–30). As well, proinflammatory cytokines secreted shortly after infection (31) may activate macrophages and enhance the phagocytosis of infected CD4^+^ T cells in proximity *in vivo*.

Putting our observation in this context, we investigated whether Vpu-mediated CD47 downregulation would facilitate macrophage infection by promoting phagocytosis of HIV-1-infected CD4^+^ T cells. In the current study, we report that Vpu downregulates CD47 from the surface of infected CD4^+^ T cells. We also show that CD47 modulation by Vpu promotes enhanced capture and phagocytosis of T cells by monocyte-derived macrophages (MDMs), which ultimately leads to productive infection of MDMs. In addition, our findings uncover that through this process a T/F virus could efficiently infect MDMs, revealing a possible model for macrophage infection at early stages of infection. Importantly, mechanistic studies reveal that Vpu depletes CD47 via a process that requires its transmembrane domain (TMD) for binding CD47 as well as the DSGNES diserine motif and the ExxxLV trafficking motif for targeting CD47 to lysosome-dependent degradation.

## RESULTS

### Vpu downregulates CD47 from the surface of HIV-1-infected CD4^+^ T cells

We and others previously identified CD47 as a putative target for Vpu-mediated surface downregulation (6, 10). To verify the expression profile of CD47 in the context of HIV-1 infection, we first examined the effect of Vpu on CD47 surface expression levels in HIV-1-infected SupT1 cells that do not express BST2 (32). Given that both BST2 and CD47 localize to lipid rafts at the cell surface (12, 33), and as such might be part of supramolecular protein complexes, the use of SupT1 cells would allow us to determine whether the effect of Vpu on CD47 was independent from BST2 downmodulation by Vpu. To this end, SupT1 cells were infected with the CXCR4 (X4)-tropic GFP-marked NL4-3 HIV-1 (NL4-3) expressing (WT) or lacking Vpu (dU) and surface expression of CD47 was measured by flow cytometry at 48 hours post-infection (hpi). Infection with WT HIV-1 resulted in an ∼30% decrease in surface CD47 levels on infected cells as compared to bystander GFP^−^ cells or cells infected with dU HIV-1, suggesting that modulation of CD47 by Vpu did not involve BST2 (Fig. 1). This downregulation of CD47 was also observed to varying extents in primary CD4^+^ T cells infected with either X4-tropic NL4-3, CCR5 (R5)-tropic NL 4-3.ADA.IRES.GFP (NL 4-3 ADA) or R5-tropic T/F WITO virus expressing Vpu. Importantly, no such modulation in CD47 expression was noted in cells infected with the respective dU derivatives of these viruses (Fig. 1), further confirming a Vpu-dependent downregulation of CD47 during HIV infection of primary CD4^+^ T cells.

**Figure 1.**
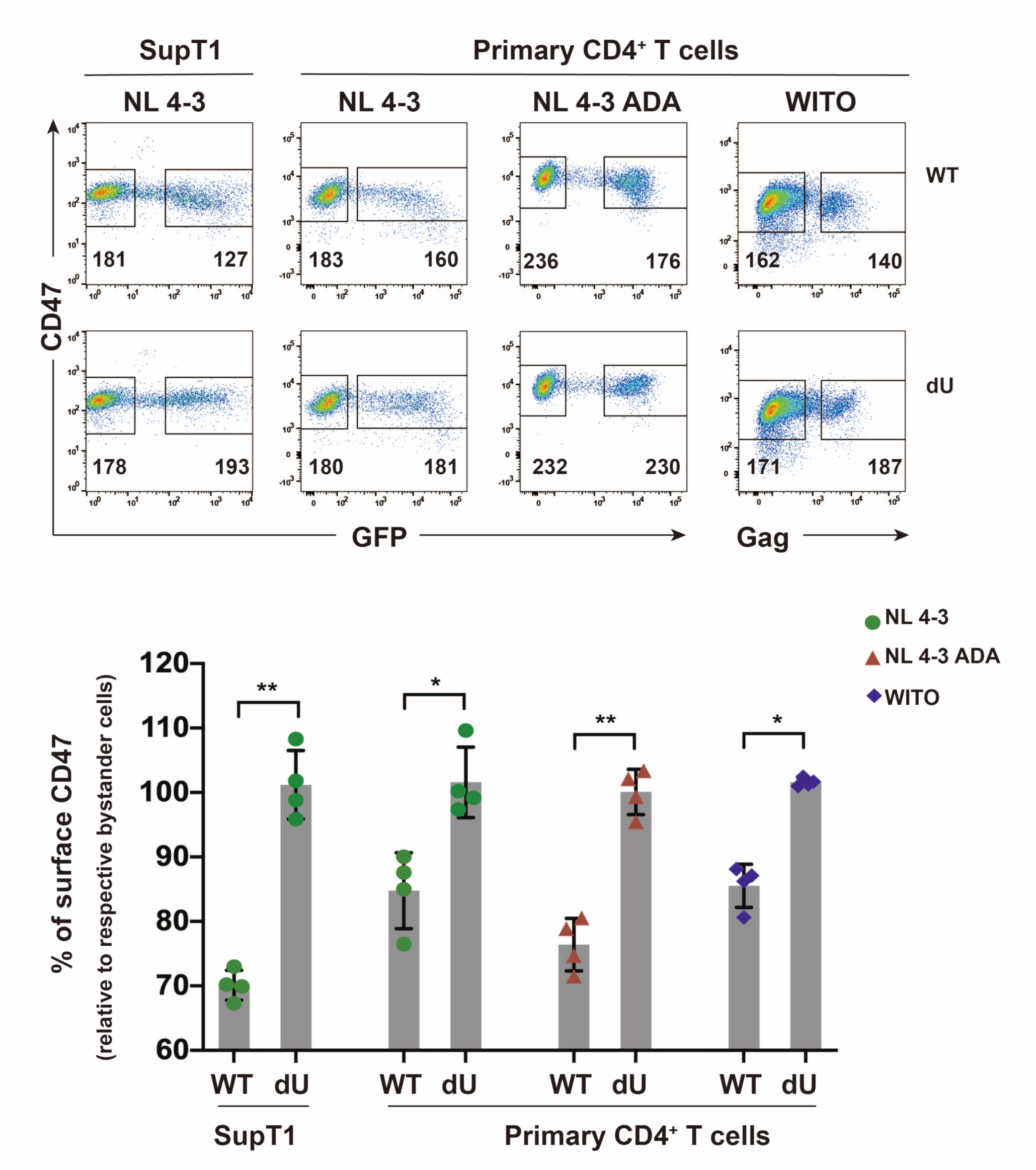
CD47 is downregulated from the surface of HIV-1-infected CD4^+^ T cells by Vpu. SupT1 T cells or primary CD4^+^ T cells were infected with GFP-expressing NL4-3 (WT or dU) viruses or with either VSV-G pseudotyped GFP-expressing NL 4-3 ADA (WT or dU) or transmitted/founder WITO (WT or dU) viruses as indicated. After 48 h, cells were stained with anti-CD47 (clone CC2C6) and analyzed by flow cytometry. **(Top)** Representative flow cytometry dot-plot graphs with indication of the median fluorescence intensity (MFI) values for infected (GFP- or Gag-positive) and bystander cells (GFP- or Gag-negative). **(Bottom)** Summary graphs of relative surface CD47 expression levels at 48 h postinfection (hpi) with the indicated viruses (n=4). The percent MFI values were calculated relative to that obtained in the respective bystander cells. Statistical analysis was performed using Mann-Whitney U-test (**, P < 0.01; *, P < 0.05), error bars represent standard deviations (SD). Flow cytometry data for this figure was generated on a CyAn ADP cytometer (Beckman coulter).

CD47 is reported to undergo post-translational pyroglutamate modification at the SIRPα binding site by the glutaminyl-peptide cyclotransferase-like protein (QPCTL), a modification that is thought to positively regulate the CD47/SIRPα axis by enhancing SIRPα binding (34). Since CD47 surface expression was detected using anti-human CD47 mAb clone CC2C6, which specifically recognizes the pyroglutamate of CD47, we next assessed whether Vpu targeting of CD47 is dependent or independent of the pyroglutamate epitope. Using the anti-human CD47 mAb clone B6H12 that recognizes all forms of CD47 at the cell surface, we found that the extent of Vpu-mediated downregulation of CD47 (∼40%) was comparable to that detected with the CC2C6 mAb in infected Jurkat E6.1 cells (Fig. S1). Together, these data show that Vpu downregulates all forms of CD47 from the surface of HIV-1-infected CD4^+^ T cells.

### Vpu-mediated CD47 downregulation enhances capture and phagocytosis of infected T cells by MDMs

CD47 is known to function as a marker of “self” that protects healthy cells from being engulfed by macrophages. Accordingly, hematopoietic cells lacking CD47 are efficiently cleared by macrophages (35). Therefore, we hypothesized that Vpu-mediated downregulation of CD47 modulates the capture and phagocytosis of HIV-1-infected T cells by MDMs. To test this, we generated a CD47 knockout (CD47KO) Jurkat E6.1 cell line (Fig. S2A and B). First, target Jurkat cells (CD47 expressing control and CD47KO) were infected with NL 4-3 ADA (WT or dU) for 48 h, then labelled with CFSE and cocultured with MDMs for 2 h to assess the capture of labelled T cells by CD11b-expressing macrophages using flow cytometry, as described in material and methods (Fig. 2A). As shown in Fig. 2B, we observed a significantly higher frequency of CD11b^+^/CFSE^+^ MDMs upon coculture with WT-infected CD47-expressing Jurkat cells (∼12 %, average of n=4) compared to those cocultured with mock- or dU-infected Jurkat cells (∼7.5 or 8.8 %, respectively, average of n=4). When MDMs were cocultured with CD47KO Jurkat cells, there was an overall increase in capture of target cells (21.5 to 23.3 %, average of n=4 for mock and WT, respectively) regardless of whether they were infected with WT or dU virus (Fig. 2B; Fig. S2C). In keeping with an inverse correlation between cell capture efficiency and CD47 expression on target cells, these results indicate that Vpu-mediated CD47 downregulation enhanced the susceptibility of T cells to be taken up by MDMs.

**Figure 2.**
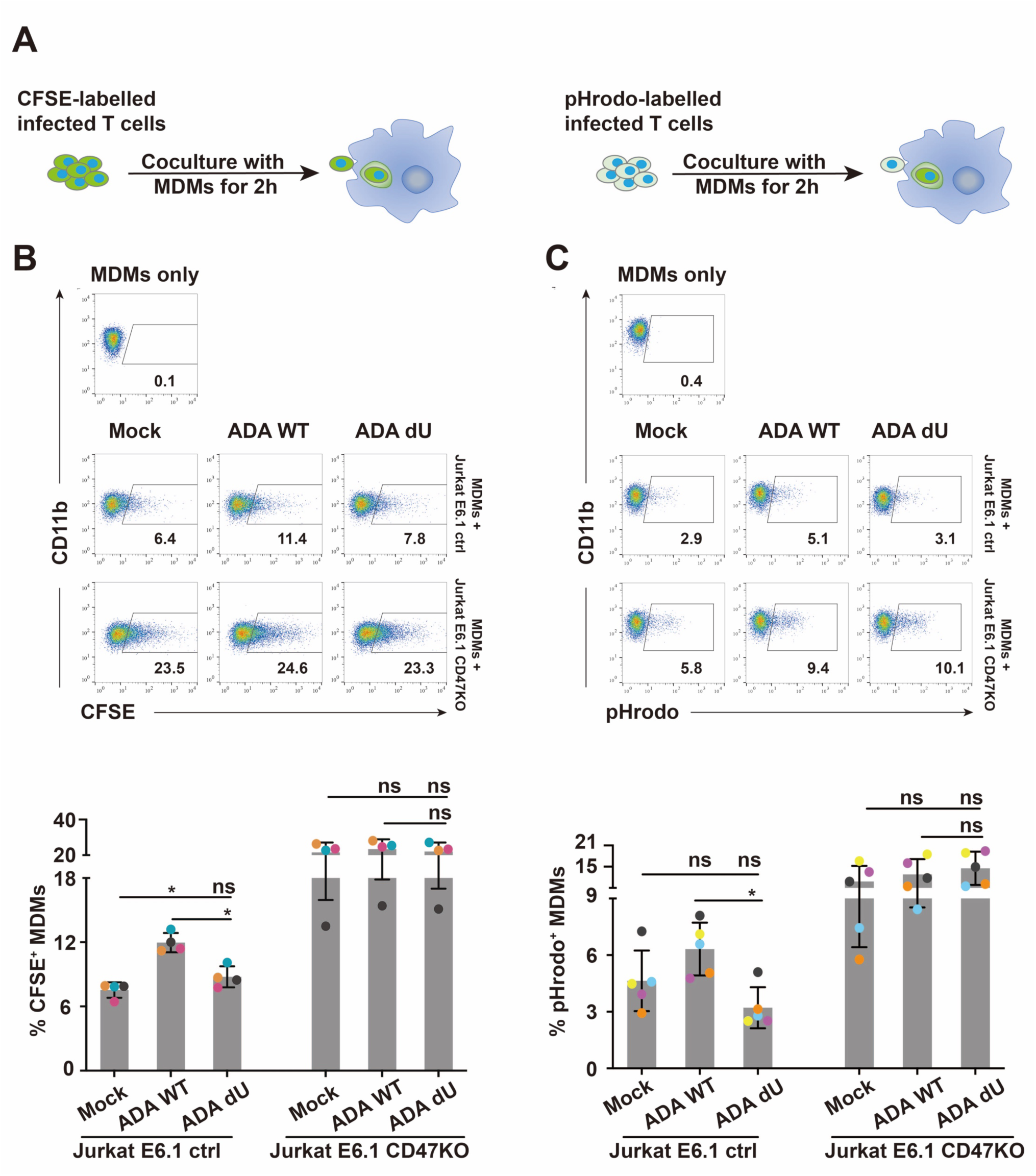
Vpu-mediated CD47 downregulation enhances capture and phagocytosis of infected T cells by MDMs. **(A)** Experimental strategy for HIV-1-infected target cells labelling, coculture with MDMs for analysis of *in vitro* capture or phagocytosis by flow cytometry. Target cells were either mock-infected or infected with VSV-G pseudotyped NL 4-3 ADA (WT or dU) viruses for 48 h and labelled with either CFSE (left) or pHrodo (right). **(B)** Representative flow cytometry dot-plots of MDMs (CD11b^+^) with percentage numbers of CFSE^+^ population corresponding to capture (top); summary graphs for capture of CFSE-labelled CD47 expressing Jurkat E6.1 control (ctrl) or CD47 knockout (KO) Jurkat E6.1 cells by MDMs in the indicated conditions (bottom). **(C)** Representative flow cytometry dot-plots of MDMs (CD11b^+^) with percentage numbers of pHrodo^+^ populations corresponding to phagocytosis (top); summary graphs for phagocytosis of pHrodo-labelled target cells by MDMs in the indicated conditions (bottom). **(B and C)** analyzed by Mann-Whitney U-test (*, P < 0.05; ns, nonsignificant, P > 0.05), error bars represent SD.

To directly demonstrate that this process was a consequence of phagocytosis, we performed similar experiments using target cells labelled with pHrodo, a pH-sensitive dye that becomes fluorescent within the acidic environment of phagolysosomes, thus enabling an accurate measurement of *bona fide* phagocytosis. Indeed, using this approach we also observed a significantly higher frequency of CD11b^+^/pHrodo^+^ cells when MDMs were cocultured with WT (6.3 %, average of n=5) instead of dU (3.2 %, average of n=5) virus-infected targets (Fig. 2C). Also consistent with the data from the capture assay (Fig. 2B; Fig. S2C), CD47KO target cells were more efficiently phagocytosed by MDMs and the extent of which was comparable among uninfected, WT- or dU-virus (10.8, 12.8 and 14.6 %, respectively, average of n=5) infected targets (Fig. 2C; Fig. S2D). Importantly the differential impact of Vpu on the capture and phagocytosis of CD47-expressing T cells was not linked to an increase in the frequency of Annexin^+^ apoptotic target cells, a condition known to trigger phagocytosis by MDMs (Fig. S2E). Taken together, these results indicate that Vpu promotes both capture and phagocytosis of target cells.

### Phagocytosis of infected CD4^+^ T cells promotes productive infection of MDMs by T/F virus

T/F viruses were reported to display a much weaker tropism for MDMs compared to truly M-tropic virus strains (27). Indeed, there was no detectable infection (based on intracellular Gag p24) of MDMs using WITO T/F virus (MOI = 5), whereas infection with a cell-free WT NL 4-3 ADA virus (M-tropic, MOI = 2) resulted in up to 1.5 - 5 % of p24^+^ cells (Fig. S3A). To investigate whether WITO could infect MDMs via phagocytosis of infected primary CD4^+^ T cells, MDMs were either cocultured with CD4^+^ T cells infected (to a comparable level, Fig. S3B) with either ADA or WITO viruses or cultured in the presence of the corresponding virion-containing supernatants from T cell cultures prior to extensive washes to eliminate input target T cells or virions. The former was referred to as “co-culture” while the latter, was designated “cell-free” in Fig. 3. Conditioned supernatant from both “co-culture” and “cell-free” infections was collected at various time points and quantified for infectious particles using a TZM-bl cell-based luciferase reporter assay (Fig. 3A). While the media from cell-free-infected MDMs revealed a modest luciferase activity for both ADA and WITO infections, that from MDMs cocultured with infected CD4^+^ T cells showed meaningfully higher levels (Fig. 3B). Interestingly, we observed an approximately 6- to 9-fold higher viral production (day 2) for MDMs cocultured with WITO-infected CD4^+^ T cells compared to those with ADA-infected cells, despite the initially comparable infection of CD4^+^ T cells at the time of the cocultures (Fig. 3B; Fig. S3B). These results show that HIV and notably T/F viruses can productively infect MDMs through cell-to-cell contact with infected CD4^+^ T cells. To directly support the notion that capture and engulfment of infected T cells was occurring in these conditions, MDMs cocultured with T cells infected with GFP-marked viruses were processed for immunostaining and analysis by confocal microscopy. As shown in Figure 3C, the presence of GFP^+^ MDMs in close contact with T cells or containing intact GFP^+^ T cells could be observed. Furthermore, MDMs displaying a GFP signal following cell-contact showed the presence of multiple nuclei and harbored Gag p17 immunostaining in internal compartments as well as at the cell periphery, suggesting the intercellular transfer of fully mature virus particles to MDM (Figure 3D). Taken together with the capture/ phagocytosis data (Fig. 2), we assert that the improved infection of MDMs is likely a consequence of their engulfment of infected CD4^+^ T cells.

**Figure 3.**
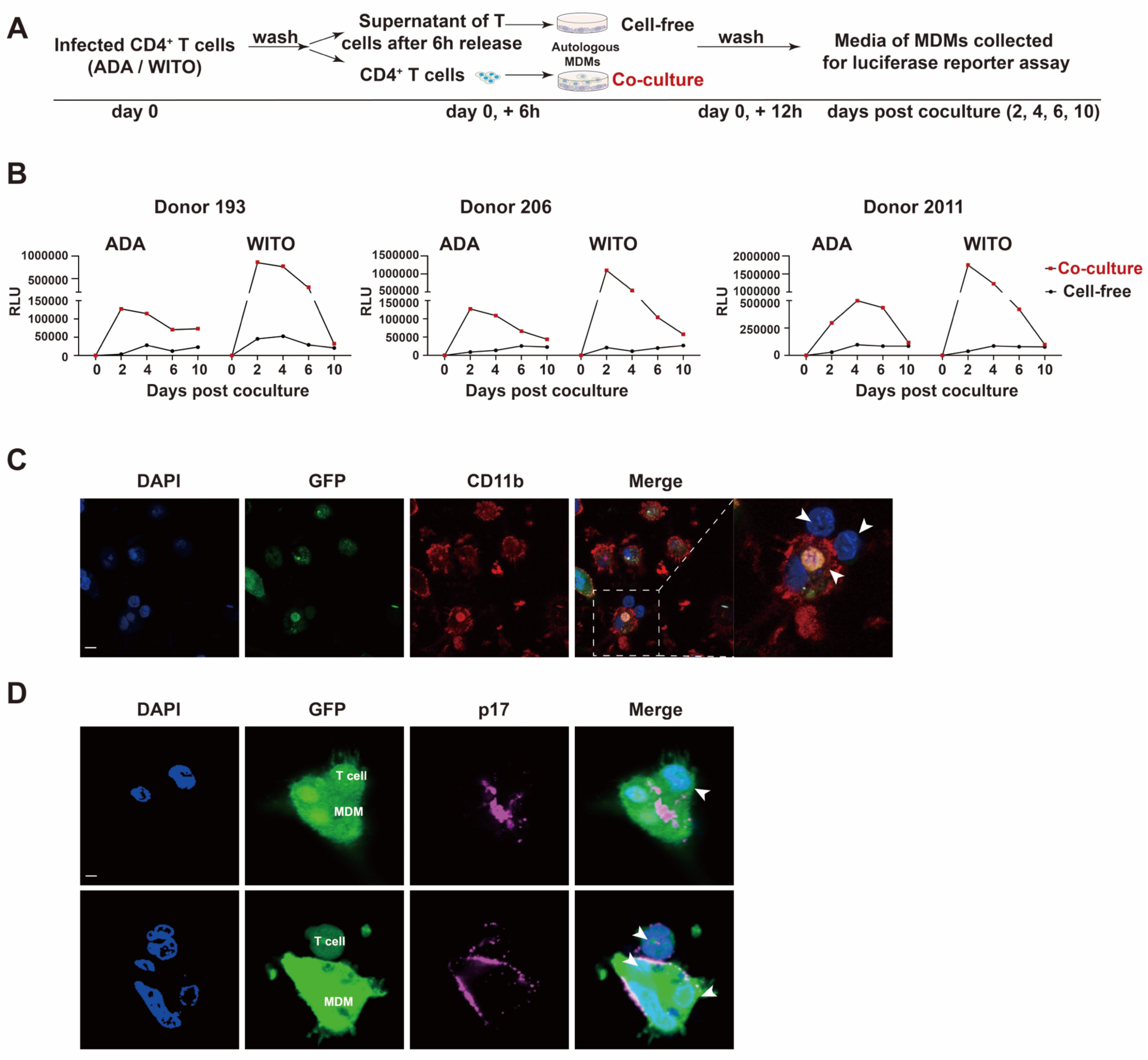
Phagocytosis of infected CD4^+^ T cells promotes productive infection of MDMs by T/F virus. **(A)** Experimental strategy for coculture infected CD4^+^ T cells with autologous MDMs, and analysis of MDMs productive infection. MDMs were cocultured for 6 h with WT NL 4-3 ADA- or WITO-infected autologous CD4^+^ T cells (**Co-culture**) or were exposed for 6 h with supernatants from the same HIV-1-infected T cells (**Cell-free**). MDMs were maintained in culture after washing-off T cells or supernatants, media of MDMs were collected at the indicated time points to assess the production of infectious particles via infection of TZM-bl cells and luciferase (Luc) activity assay. **(B)** TZM-bl cells were infected for 48 h with media of MDMs collected at different time points and assayed for Luc activity. Results are expressed as relative light units (RLU). Shown are RLU of TZM-bl infected with media collected from MDMs from 3 donors. **(C-D)** GFP-expressing NL 4-3 ADA-infected SupT1 cells were cocultured for 2 h with MDMs, cells were then stained with anti-CD11b (C) or anti-p17 Abs (D) as well as DAPI and analyzed by confocal microscopy (scale bar, 10 µm), T cells are indicated by white arrows.

To provide evidence that the infection of MDMs was indeed due to phagocytosis, we introduced Jasplakinolide (Jasp) to the cocultures (Fig. 4A). Jasp was reported to promote actin polymerization and stabilize actin filaments, thereby inhibiting cellular processes dependent on actin dynamics including phagocytosis (28, 36). In brief, MDMs were pretreated with Jasp, cocultured with WITO-infected CD4^+^ T cells, and analyzed by flow cytometry for phagocytosis activity (Fig. 4B) and the frequency of infected (Fig. 4C) MDMs. Treatment of MDMs with Jasp effectively inhibited phagocytosis (Fig. 4B) and blocked the infection (Fig. 4C). Consistent with these findings, the level of infectious viral particles, measured via luciferase activity in TZM-bl cells was negligible in the supernatant of MDMs cultures following Jasp treatment (Fig. 4D). As well, the fact that reverse transcriptase inhibitor zidovudine (AZT) or integrase inhibitor raltegravir (Ral) (Fig. S4A and B) could block MDMs infection following coculture of WITO-infected CD4^+^ T cells further validates the authenticity of the MDMs infection.

**Figure 4.**
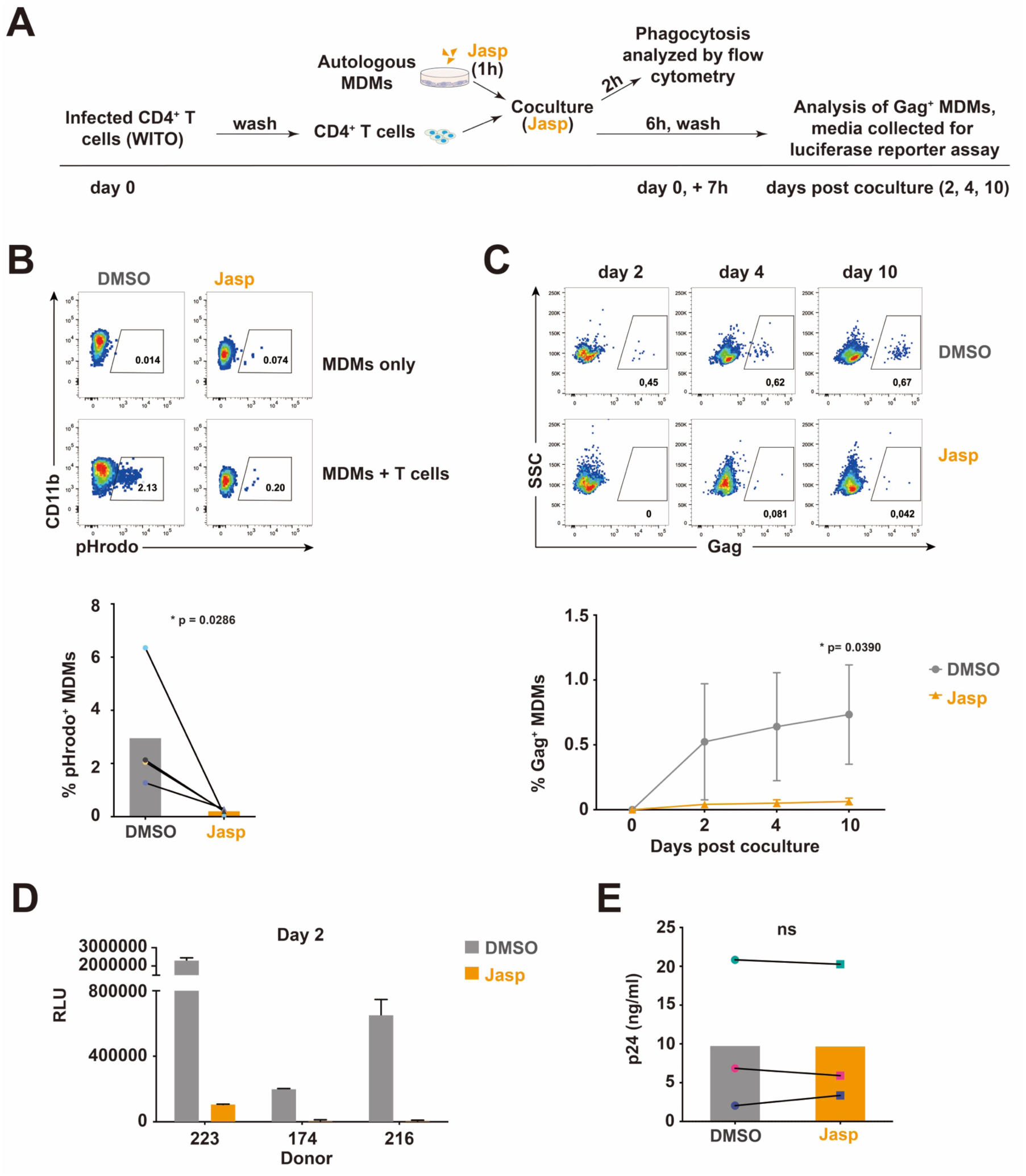
Inhibition of phagocytosis hinders productive infection of MDMs by T/F virus. **(A)** Experimental strategy for coculture of infected CD4^+^ T cells, with autologous MDMs pretreated with Jasplakinolide (Jasp), analysis of phagocytosis and MDM productive infection. Pretreated MDMs (1 h with Jasp or vehicle (DMSO)) were cocultured for 6 h with WITO-infected CD4^+^ T cells in the presence of Jasp or DMSO. MDMs were also cocultured for 2 h with the same number of pHrodo-treated CD4^+^ T cells and analyzed for phagocytosis by flow cytometry. MDMs were maintained in culture after washing-off T cells and collected at the indicated time points for intracellular Gag staining and flow cytometry analysis. Evaluation of infectious virus production was determined as described above using the TZM-bl assay. **(B)** Inhibition of phagocytosis by Jasp. Representative flow cytometry dot-plots of MDMs (CD11b^+^) with percentage of pHrodo^+^ populations corresponding to phagocytosis of CD4^+^ T cells by MDMs (top) and summary graph (bottom), analyzed by Mann-Whitney U-test, (*, P < 0.05). **(C)** Inhibition of MDMs infection by Jasp. Representative flow cytometry dot-plots of MDMs showing the percentage of Gag^+^ cells at the indicated time points (top) following exposure to infected target cells; summary graph represents the data obtained with MDMs of 3 distinct donors (bottom). **(D)** TZM-bl cells infected with media from MDMs cocultured with infected CD4^+^ T cells were assayed for Luc activity; shown are RLU detected with media collected from MDMs from 3 distinct donors. **(E)** Jasp does not affect viral release from infected CD4^+^ T cells. WT WITO virus-infected cells were washed to remove cell-associated virions, and then treated with Jasp or vehicle DMSO for 6 h. Virus-containing supernatants were collected and quantified for p24 by ELISA; shown are the results obtained with CD4^+^ T cells from 3 distinct donors. **(C and E)** Analyzed by two-tailed Student’s *t*-test, (*, P < 0.05; ns, nonsignificant, P>0.05), error bars represent SD.

Given that the effect of actin filament disruption on HIV-1 viral release remained unclear (37), we assessed whether Jasp treatment would affect virion release, which would ultimately interfere with MDMs infection. To this end, WITO-infected T cells were washed to remove cell-free virions and then cultured in the presence or absence of Jasp. As shown in Fig. 4E, Jasp treatment did not affect the release of viral particles from T cells. Altogether, using different cell-based assays, we provide evidence that the ability of MDMs to phagocytose infected T cells rendered them susceptible to infection by viruses which would otherwise be poorly infectious.

### Vpu binds CD47 via its transmembrane domain (TMD) and targets CD47 for lysosomal degradation

We next sought to understand the mechanism involved in Vpu-mediated CD47 antagonism. HEK 293T cells were co-transfected with plasmids expressing CD47 and Vpu and analyzed by Western blotting for CD47 expression. CD47 was downregulated by Vpu in a dose-dependent manner by as much as 70% (Fig. 5A), bringing the question as to how, mechanistically, Vpu mediates the depletion. Thus, we generated Vpu variants that contain mutations within the main functional domains including the: (1) A15L-W23A in the TMD that is involved in various target interactions (38, 39), (2) S53/57A mutation within the DSGNES diserine motif that is involved in the recruitment of the SCF^βTrCP^ E3 ubiquitin ligase, responsible for ubiquitination and degradation of several Vpu targets (40, 41), and (3) A_63_xxxA_67_V (AxxxAV for short) within the ExxxLV trafficking motif, which targets Vpu-containing complexes to intracellular compartments away from the plasma membrane (42). To this end, we found that all three Vpu mutants prevented CD47 depletion (Fig. 5B and C(input)), suggesting the importance of the main functional domains of Vpu in this process. Indeed, the A15L-W23A mutant was unable to bind CD47 (Fig. 5C), implying that the Vpu TMD mediates the complex formation with CD47. To determine whether Vpu induces CD47 protein degradation by either the proteasomal or lysosomal pathway, we treated Vpu- and CD47-expressing HEK 293T transfectants with proteasomal inhibitor MG132 or lysosomal inhibitor Concanamycin A (ConA) and found that ConA, but not MG132 prevented CD47 depletion (Fig. 5D). Collectively, these results indicate that Vpu binds CD47 via its TMD and targets the host protein for lysosomal degradation. Conceivably, this process requires both the SCF^βTrCP^-recruiting DSGNES diserine motif as well as the ExxxLV trafficking signal, consistent with the lack of CD47 degradation by the S53/57A and AxxxAV mutants.

**Figure 5.**
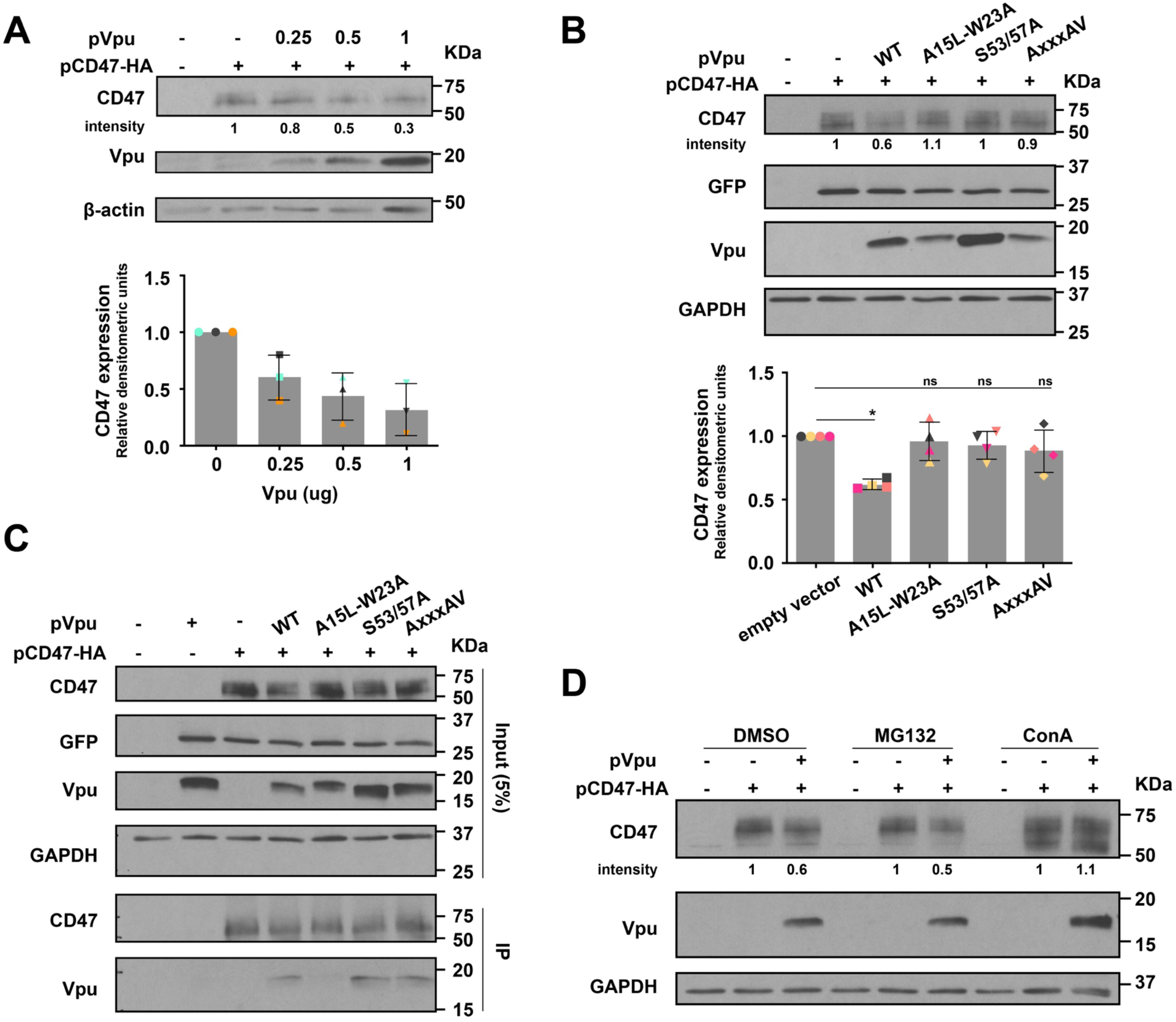
Vpu binds CD47 via its transmembrane domain (TMD) and targets CD47 for lysosomal degradation. **(A)** Vpu induces depletion of CD47. HEK 293T cells were cotransfected with plasmids encoding HA-tagged CD47 (pCD47-HA), along with increasing concentrations of GFP-marked plasmids expressing wild-type ADA Vpu (pVpu). An empty vector expressing GFP alone was added to adjust the total amounts of plasmid DNAs in all conditions. Whole cell lysates were analyzed for the indicated proteins by Western blotting. A Representative blot is shown (top), and a summary graph of densitometric analysis of CD47 is presented (bottom), error bars represent SD. **(B)** Vpu-mediated CD47 depletion requires the main Vpu functional motifs. HEK 293T were cotransfected with pCD47-HA, along with either empty vector, or plasmids encoding WT Vpu, or the indicated Vpu mutants. A representative Western blot is shown (top) as well as a summary graph of densitometric analysis of CD47 (bottom); statistical significance was determined by Mann-Whitney U-test (*, P < 0.05; ns, nonsignificant, P > 0.05); error bars represent SD. **(C)** HEK 293T cells were co-transfected with the indicated plasmids for 48 h prior to cell lysis and immunoprecipitation (IP) using anti-HA antibody. The immunoprecipitates were analyzed for the indicated proteins by Western blotting. **(D)** HEK 293T cells were cotransfected with the indicated plasmids for 36 h and vehicle (DMSO), MG132 or Concanamycin A (ConA) were added 8 h before cells were harvested and analyzed by Western blotting.

Furthermore, given that Vpu is typically involved in TMD-TMD interactions with its target proteins, we generated a chimeric CD47 mutant, composed of the extracellular domain (ECD1, aa 1-141) of human CD47 as well as the five membrane-spanning domains (MSDs) and the cytoplasmic tail (CT) of mouse CD47 (Fig. S5), which displays ∼26% of aa sequence divergence mainly found in the first and second MSDs. In this configuration, CD47 became largely resistant to Vpu-mediated degradation, consistent with the fact that the mouse CD47 counterpart was unsensitive to Vpu (Fig. S5). These results suggest that the MSDs are important determinants of human CD47 susceptibility to Vpu-mediated degradation.

### HIV-1-infected cells expressing Vpu-resistant chimeric CD47 are less prone to infect macrophages through phagocytosis

Given that chimeric CD47 was resistant to Vpu-mediated degradation, we next asked if expression of this mutant would alter target cell susceptibility to phagocytosis by MDMs. To this end, we used a CD47KO Jurkat cell line (JC47) (43) to generate cell lines stably expressing either the human-mouse chimeric CD47 (JC47-cCD47) or human CD47 (JC47-hCD47). Cells were selected and enriched by fluorescence-activated cell sorting (FACS) to obtain CD47 expression levels comparable to those detected on the parental Jurkat cells (Fig. 6A). Upon infection of these cell lines with NL 4-3 ADA WT HIV-1, we observed downregulation of CD47 by 40 % on JC47-hCD47 cells but only by 10 % on those expressing the chimeric JC47-cCD47 (Fig. 6B). Next, we investigated the susceptibility of uninfected (mock) and infected JC47-derived cell lines to phagocytosis by MDMs. First, we found that JC47-hCD47 cells were phagocytosed by MDMs to a similar degree as JC47-cCD47 cells, since both showed ∼ 5% CD11b^+^/pHrodo^+^ cells, suggesting that cCD47 was as effective as hCD47 at inducing a “don’t-eat-me” signal (Fig. 6C and Fig. 2C). Interestingly, upon infection with WT HIV-1, JC47-hCD47 cells were opsonized more efficiently than infected JC47-cCD47 cells (Fig. 6C). Importantly, this difference in phagocytosis was not linked to apoptosis of target cells (Fig. S6). Furthermore, and in agreement with the phagocytosis results, we observed a heightened virus production as measured by luciferase activity from MDMs co-cultured with WT HIV-infected JC47-hCD47 cells (Fig. 6D) compared to their chimeric JC47-cCD47 counterparts. Collectively, these results further underscore our observation that Vpu-mediated CD47 downregulation potentiates phagocytosis of infected T cells by MDMs and, consequently, promotes increased productive infection of MDMs.

**Figure 6.**
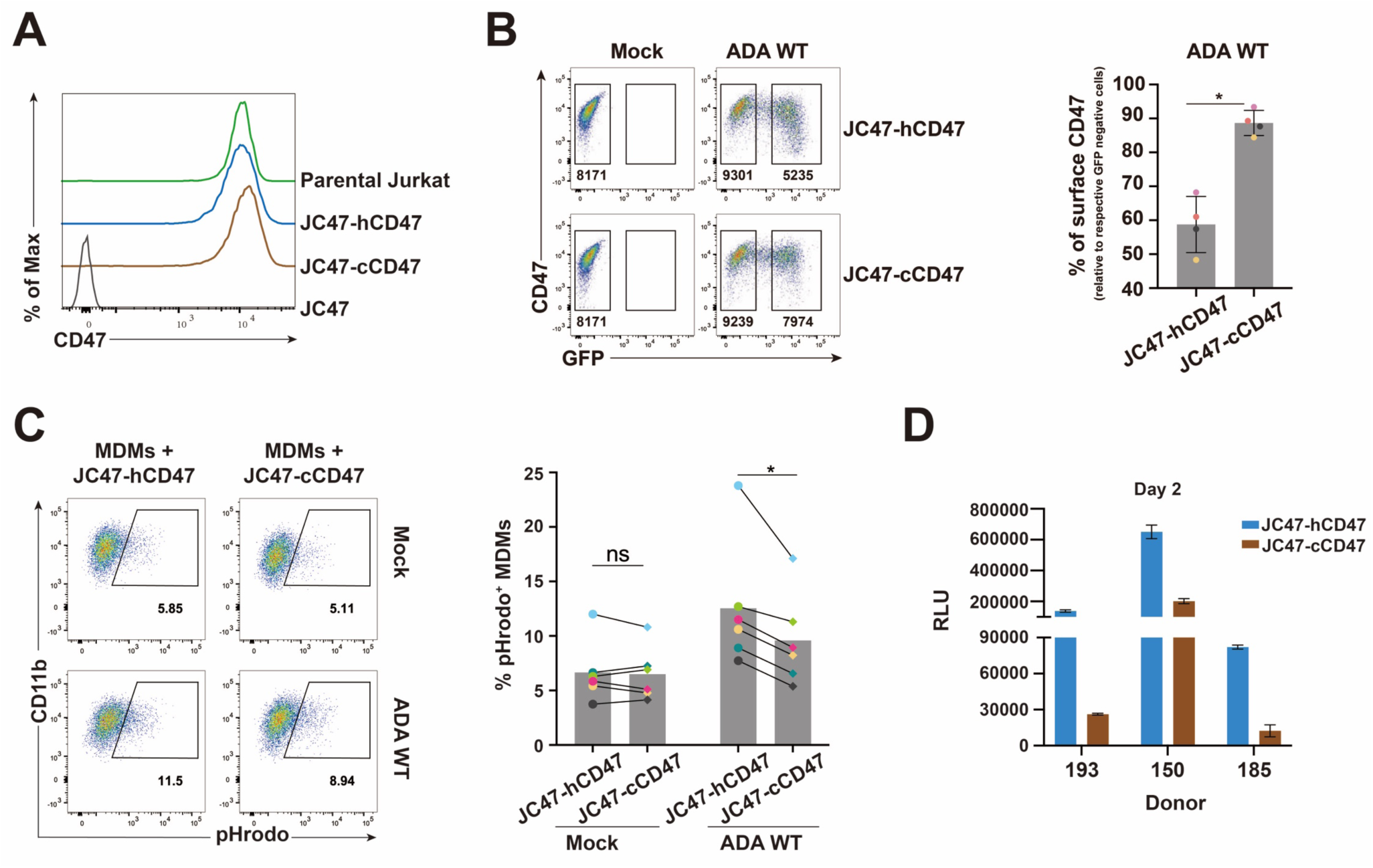
HIV-1-infected CD4^+^ T cells expressing Vpu-resistant chimeric CD47 are less prone to infect macrophages through phagocytosis. **(A)** Flow cytometry histogram to validate CD47 surface expression levels in different target Jurkat cell lines including JC47 (CD47 knockout), JC47-hCD47 (human CD47 reintroduced in JC47), JC47-cCD47 (chimeric CD47 reintroduced in JC47). **(B)** JC47-hCD47 and JC47-cCD47 cells were mock-infected or infected with VSV-G pseudotyped GFP-expressing WT NL 4-3 ADA virus for 48 h, then stained with anti-CD47mAb (clone CC2C6) and analysed by flow cytometry. Representative flow cytometry dot-plot graphs with the MFI values in infected (GFP-positive) and bystander cells (GFP-negative), (left); summary graphs of relative surface CD47 expression levels at 48 h after infection (n=4), (right), the percent MFI values were calculated relative to respective their GFP-negative cells. Statistical analysis was performed using Mann-Whitney U-test (*, P < 0.05); error bars represent SD. **(C)** The indicated mock or HIV-1-infected target cells were labelled with pHrodo and cocultured with MDMs for 2 h, prior to analysis of MDMs by flow cytometry. Representative flow cytometry dot-plots of MDMs (CD11b^+^) with percentage of pHrodo^+^ populations corresponding to phagocytosis of target cells by MDMs (left); summary graphs for MDMs (right) from 6 distinct donors, analyzed by Wilcoxon matched-pairs signed rank test (*, P < 0.05; ns, nonsignificant, P > 0.05). **(D)** JC47-hCD47 or JC47-cCD47 cells were infected as described and cocultured with MDMs for 6 h. Following washing-off of infected T cells, MDMs were cultured for 2 days, media of MDMs was collected to infect TZM-bl, for luciferase assay. Shown are RLU detected with media collected from MDMs of 3 distinct donors, error bars represent SD.

## DISCUSSION

In this study, we extend our previous SILAC-based observation that CD47 is a putative target of HIV-1 Vpu (6, 10) and reveal that Vpu indeed downregulates CD47 on CD4^+^ T cells infected with lab-adapted X4-tropic NL 4-3, R5-tropic NL 4-3 ADA as well as T/F WITO virus (Fig. 1). These findings obtained in the context of HIV-1 infection are in contrast to those reported by the Hasenkrug group, which showed that CD47 was upregulated in different types of immune cells upon recognition of several pathogens including SARS-CoV-2, hepatitis C virus (HCV) and lymphocytic choriomeningitis virus (LCMV) (44, 45), suggesting that downregulation of CD47 is an evolved viral counter-measure which provides HIV with a selective advantage. Indeed, we show that Vpu-mediated CD47 downregulation leads to an enhanced capture and phagocytosis of infected CD4^+^ T cells by MDMs, a process which ultimately leads to productive infection of macrophages. As well, our data show that the R5-tropic WITO T/F virus relies on phagocytosis to efficiently infect MDMs, raising the possibility that phagocytosis of infected cells is an important mechanism through which myeloid cells get productively infected by HIV-1 and is likely consequential for inter-host HIV-1 transmission.

Macrophages were reported to engulf HIV-1-infected CD4^+^ T cells, a process that leads to their own infection (28, 29), but the capture recognition signals remained unclear. Binding of CD47 to SIRPα suppresses multiple pro-phagocytosis signaling pathways including those mediated by IgG/FcγR, complement/complement receptors, and calreticulin (46, 47), suggesting that decreased CD47 expression might trigger phagocytosis. Indeed, we show that Vpu-mediated CD47 downregulation enhanced capture and engulfment of infected T cells by MDMs (Fig. 2). Although modest, the effect was invariably reproducible and consistent with other studies which used Jurkat cell lines expressing differential surface levels of CD47 as target cells in phagocytosis assays (43, 48). Apart from calreticulin, phosphatidylserine (PS) is another critical pro-phagocytosis signal predominant on apoptotic cells (49). Nevertheless, CD47/ SIRPα signaling was recently reported to block the “eat-me” signal driven by PS (50). In the context of HIV-1 infection, we found more externalization of PS on the cell surface as measured by Annexin V staining (Fig. S2E), in line with previously reported results (51). However, although Vpu expression was reported to induce apoptosis (52), we did not observe a significant difference of PS exposure between cells infected with WT and dU NL 4-3 ADA HIV-1 (Fig. S2E) suggesting that the augmented phagocytosis of cells infected by WT virus was likely resulting from Vpu-induced decrease in CD47. Indeed, this notion was further supported by our finding that target T cells expressing human-mouse chimeric form of CD47 that are less responsive to Vpu-mediated downregulation were less prone to phagocytosis by MDMs as compared to human CD47 expressing target cells (Fig. 6C). Furthermore, since we show that knock-out of CD47 in Jurkat T cells results in a strong enhancement of phagocytosis by MDMs (Fig. S2D), our data are collectively consistent with findings showing that lack of CD47 results in augmented opsonization of red blood cells (35) and more efficient clearance of lymphohematopoietic cells by macrophages (53). Conversely, target cells with elevated cell surface CD47 are shown to be protected from phagocytosis by macrophages (48, 54).

Infection of macrophages can be detected throughout all stages of HIV-1 infection (26). However, M-tropic viruses are found at late stages of infection and virus isolated at early stages of infection display very limited tropism for macrophages (17, 25). We hypothesized that through phagocytosis of T/F virus-infected CD4^+^ T cells, macrophages could become infected with these viruses, a process that would potentially initiate inter-host viral dissemination at early stages of HIV-1 infection. Indeed, we found that T/F virus WITO, which poorly infects MDMs by a cell-free route (Fig. S3A), as reported previously (27, 28, 55), was able to elicit a productive infection in MDMs. Interestingly, we observed that phagocytosis of WITO-infected CD4^+^ T cells ultimately led to an approximately 6- to 9-fold higher viral production by infected MDMs compared to the engulfment of T cells infected with a M-tropic NL 4-3 ADA (Fig. 3B) despite comparable degree of CD47 downregulation by these viruses (Fig. 1). This implies a potentially more efficient infection of MDMs following opsonization of T cells infected with T/F WITO since the frequency of infected T cells was comparable at the time of phagocytosis (Fig. S3B).This difference in infection efficiency between T/F WITO and HIV ADA may be linked to the differential impact of host antiviral restriction factors, and notably interferon-induced transmembrane proteins (IFTIMs), which accumulate intracellularly during HIV-1 infection of macrophages (56, 57) and are incorporated into virions, thus reducing their infectivity (57). In this context, it was reported previously that as IFITMs incorporation increased (57), there was a decrease in the infectivity of virions produced by HIV-1 ADA-infected MDMs. In contrast, T/F viruses were reported to be relatively resistant to IFITMs-mediated restriction (58, 59) with WITO displaying the most resistance to IFITM3 and releasing the highest levels of infectious virions among all viral strains tested (59). That being said, given the gradual decrease in the level of infectious particles released by MDMs (Fig. 3B), it is possible that other restriction factors present in macrophages including GBP5 (60) and MARCH8 (61), which inhibit the infectivity of macrophage-derived virions, could play a role in controlling viral dissemination.

Phagocytosis is a process known to be important for the elimination of engulfed pathogens and apoptotic cells (62). However, we show that inhibition of phagocytosis by Jasp is linked to a suppression of productive infection of MDMs (Fig. 4), suggesting that HIV-1 takes advantage of this process to infect macrophages. Indeed, a recent study by the Kieffer group (29) in HIV-1-infected humanized mice provided evidence that bone marrow macrophages phagocytosed infected T cells and produced virus within enclosed intracellular compartments. Using electron tomography (ET), they observed macrophages phagocytosing infected T cells, with mature and immature HIV-1 virions within macrophage phagosomes alongside engulfed cells at varying degrees of degradation. Moreover, virions were also observed to assemble and undergo budding and maturation within fully-enclosed compartments which would subsequently fuse with surface-accessible invaginations to release virions into the extracellular space. Since HIV-1 virions are inactivated in acidic environments (63), further investigation is needed to better understand how virions escape phagosomal degradation before a complete destruction of the ingested T cells. In fact, many microorganisms have evolved multiple strategies to prevent phagocytic destruction. For instance, *Mycobacterium tuberculosis* inhibits the acidification process of phagosomes via the exclusion of vesicular proton-ATPase thus hindering the maturation of these compartments (64); it also prevents the fusion of lysosomes with phagosome (65). Interestingly, we show herein that inhibition of reverse transcription and integration suppresses productive infection of MDMs by WITO (Fig. S4), suggesting that virions transferred to macrophages via phagocytosis of HIV-1-infected T cells were able to actively replicate in MDMs.

While phagocytosis of infected T cells by macrophages represents one route of infection of macrophages, other mode of cell-to-cell virus transfer have also been described in vitro. It was recently reported that contacts between infected T lymphocytes and macrophages could lead to virus spreading to macrophages via a two-step fusion process that involves fusion of infected T cell to macrophages and virus transfer to these newly formed lymphocytes/macrophages fused cells. These newly formed cells were in turn able to fuse to neighboring uninfected macrophages leading to the formation of long-lived virus-producing multinucleated giant cells (MGCs) (66, 67). Although the formation of MGCs has been reported in the lymphoid organs and central nervous system of HIV-1-infected patients (68–70) and SIV-infected macaques (71), the presence of macrophage-T-cell fusion was not observed by the Kieffer group in humanized mice (29).

In the context of experiments aimed at investigating whether phagocytosis of infected T cells ultimately leads to productive infection of macrophages, we have intentionally avoided the comparison between WT or dU-infected target T cells. Considering that virions are tethered at the cell surface by BST2 in the absence of Vpu, phagocytosis of dU-infected CD4^+^ T cells or fusion between these infected T cells and macrophages could inadvertently result in higher productive infection of MDMs. To circumvent this problem, we generated a Jurkat cell line expressing human-mouse chimeric CD47, which is relatively resistant to Vpu modulation (Fig. S5; Fig. 6A and B). Taking advantage of this system, we confirm that Vpu-mediated CD47 downregulation contributes to phagocytosis of HIV-1-infected CD4^+^ T cells by MDMs, leading to enhanced productive infection of MDMs (Fig. 6C and D).

Mechanistically, we provide evidence that the main functional domains of Vpu are involved in the downregulation of CD47 and that Vpu interacts with CD47 via its TMD to target the latter for degradation via a lysosomal pathway (Fig. 5). Although the model of CD47 degradation seems rather similar to how Vpu depletes BST2; it remains unclear whether: (1) the DSGNES motif through the recruitment of SCF^βTRCP1/2^ complex promotes CD47 ubiquitination and (2) interaction of adaptor proteins to the ExxxLV trafficking motif of Vpu in complex with CD47 target CD47 to cellular compartment away from the plasma membrane. More detailed mechanistic studies are required to fully dissect processes underlying Vpu-mediated downregulation of CD47.

In summary, we report herein that CD47 is a new cellular target downregulated by HIV-1 Vpu. Such a decrease in CD47 expression allows for enhanced phagocytosis of infected T cells by macrophages, which ultimately leads to productive infection of this myeloid cell subset even with HIV strains that would otherwise be weakly M-tropic (i.e., T/F viruses). We posit that this process enables macrophages to be infected, including during early stages of HIV infection when M-tropic strains have not yet emerged. Taken together, our data identifies a mechanism whereby T/F virus-infected macrophages could be a source of viral reservoirs and promote viral dissemination to different tissues.

## MATERIALS AND METHODS

### Antibodies

For flow cytometry, the following antibodies (Abs) were used: PE/Cy7-conjugated mouse anti-human CD47 (clone CC2C6) monoclonal Ab (mAb) and APC-conjugated anti-CD11b mAb (clone ICRF44) as well as corresponding isotype controls from BioLegend, APC-conjugated mouse anti-human CD47 mAb (clone B6H12; eBioscience), RD1-conjugated anti-Gag (clone KC57; Beckman Coulter). For immunoprecipitation and Western blot analysis, the following Abs were used: polyclonal sheep anti-human CD47 (AF4670) and sheep IgG HRP-conjugated Ab (HAF016) from R&D system; mouse anti-HA mAb (16B12), anti-GAPDH (FF26A/F9) and anti-CRISPR Cas9 (7A9) from BioLegend; rabbit anti-HA mAb (C29F4) from Cell Signaling Technology; rabbit anti-GFP (SAB4301138) from Sigma-Aldrich; anti-β-actin (C4, sc-47778) from Santa Cruz Biotechnology; goat anti-rabbit IgG H+L (HRP, ab205718) and goat anti-mouse IgG H+L (HRP, ab 205719) from Abcam and anti-Vpu rabbit polyclonal serum as described previously (72). For confocal microscopy analysis, the following Abs were used: purified anti-CD11b (clone ICRF44; BioLegend), anti-p17 as previously described (73), Alexa Fluor 594-coupled donkey anti-mouse IgG H+L (Invitrogen, #A-21203).

### Plasmids

The X4-tropic proviral construct pBR NL 4-3. IRES. GFP wild-type (WT) and its Vpu-deficient derivative (dU) were kindly provided by Frank Kirchhoff (74, 75). The R5-tropic pNL 4-3 ADA. IRES.GFP WT and dU were generated as described (76). The molecular clone of T/F virus WITO was obtained from the NIH AIDS Reagent Program (#11919)(55) and the dU version of WITO was generated by overlapping PCR.

The pSVCMV-VSV-G plasmid encoding for the vesicular stomatitis virus glycoprotein G (VSV-G) was previously described (32). The lentiviral psPAX2 packaging vector was provided by Didier Trono (Addgene plasmid #12260). The lentivectors lentiCRISPR v2 (plasmid #52961) (77) and pWPI-IRES-Puro-Ak (plasmid #154984) were also obtained through Addgene from Feng Zhang and Sonja Best, respectively.

Vpu mutants were generated using PCR-based Quick-change site-directed mutagenesis as per standard protocols (Agilent).The plasmids encoding WT Vpu and Vpu mutants were generated by insertion of the corresponding Vpu fragments from pNL 4-3 ADA proviral constructs into the pCGCG-IRES-GFP plasmid, a kind gift from Frank Kirchhoff (75). The cDNA of human CD47 and mouse CD47 with an HA-tag at the C-terminal were purchased from Sino Biological and Thermo Fisher Scientific, respectively. The HA-tag was then added to human CD47 by PCR. Chimeric CD47 consisting of a human extracellular domain and mouse MSDs+ HA-tagged cytosolic tail was generated by overlapping PCR (see Supplementary table 1 for oligonucleotides). These fragments were then inserted into pECFP-N1 (Clonetech). Fragments without the HA-tag were also generated by PCR and inserted into pWPI-IRES-Puro-Ak to create pWPI-hCD47 or pWPI-cCD47 for expression of human or chimeric CD47, respectively. All constructs were confirmed by sequencing.

### Cell lines

HEK 293T cells and the HeLa TZM-bl indicator cell line were cultured in DMEM (Wisent) containing 100 U/mL penicillin, 100 mg/mL streptomycin (P/S) and 10% FBS (DMEM-10). Lymphocytic cell lines were maintained in RPMI 1640 medium (Wisent) containing P/S and 10% FBS (RPMI-10). SupT1 (Dr. Dharam Ablashi (78)) and TZM-bl (Dr. John C. Kappes, and Dr. Xiaoyun Wu (79)) cells were obtained from the NIH AIDS Reagent Program, whereas Jurkat E6.1 and HEK 293T cells were acquired from ATCC. The Jurkat E6.1-based CD47 knockout (KO) cell line, JC47, was as described previously (43).

### CD47 knockout and rescue

To generate a CD47KO Jurkat E6.1 cell line, guide sequence 5’ CACCGGATAGCCTATATCCTCGCTG-3’ targeting CD47 was inserted into the lentiCRISPR v2 vector. Lentiviruses were produced by triple transfection of the generated lentivector with psPAX2 and pSVCMV-VSV-G in HEK 293T cells as described previously (6). Control lentiviruses were also produced using the lentiCRISPR v2 without sgRNA. Jurkat E6.1 cells were transduced with either the control or sgRNA-expressing lentiviruses, selected with puromycin and the CD47KO population was enriched by fluorescence-activated cell sorting (FACS).

In order to rescue CD47 expression in CD47 KO cells, the JC47 cell line was used. Lentiviruses were produced by transfecting HEK 293T cells with pWPI-hCD47 or pWPI-cCD47 and psPAX2 and pSVCMV-VSV-G. JC47 cells were transduced with lentiviruses expressing either hCD47 or cCD47, selected by CD47 surface expression and enriched by FACS to ensure that CD47 expression levels are comparable with parental Jurkat E6.1.

### Primary cell cultures

Human blood samples were obtained from healthy adult donors following informed consent in accordance with the Declaration of Helsinki under research protocol approved by the Research Ethics Review Board of the Institut de Recherches Cliniques de Montréal (IRCM). Peripheral blood mononuclear cells (PBMCs) were purified from buffy coats following Ficoll density gradient (GE Healthcare). CD4^+^ T cells were isolated by negative selection using CD4^+^ T cell isolation kit (Miltenyi Biotec) according to the manufacturer’s protocol. Purified CD4^+^ T cells were activated with 5 µg/mL phytohemagglutinin-L (PHA-L; Sigma-Aldrich, #11249738001) and 100 U/mL IL-2 (PeproTech, #200-02) for 3 days and cultured in RPMI-10 containing 100 U/mL IL-2 for another 2 days before infection.

PBMCs were seeded for 2 h at 37°C in non-tissue culture treated dishes (Fisherbrand) containing serum-free RPMI medium. After gentle washes, adherent cells (which mostly contain monocytes) were cultured for 7 days in RPMI supplemented with 5% decomplemented autologous human blood plasma and 10 ng/mL M-CSF (R&D system, #216-MC) to obtain MDMs. Purity of MDMs was determined by CD11b surface staining and was found to routinely reach > 95 %.

### Virus production and infection

Virus stocks were obtained by transfecting of HEK 293T cells with proviral DNA in the presence or absence of pSVCMV-VSV-G using polyethylenimine (PEI; Polyscience, #23966). Briefly, HEK 293T cells were plated at 5 x 10^6^ cells per 15 cm dish for overnight incubation and then transfected with 20 µg of total DNA combined with 60 µg of PEI. Media was changed at 18 h post-transfection. Virus-containing supernatants were collected at 48 h post-transfection, clarified and pelleted by ultracentrifugation onto a 20% sucrose-phosphate-buffered saline (PBS) cushion for 2 h at 35,000 rpm at 4°C. Viruses were titrated using the TZM-bl indicator cells as previously described (80).

For infection of T cell lines, cells were infected at multiplicity of infection (MOI) of 0.5 or 1. Primary CD4^+^ T cells were infected at MOI of 1 by spin-inoculation as previously described (81). MDMs (seeded at 1 x 10^5^ cells/well in 12 well-plate) were infected at MOI of 2 (for NL 4-3 ADA) or MOI of 5 (for WITO) in 300 µL of RPMI-10. Viruses were adsorbed for 6 h at 37°C before medium was replaced with 1 mL of RPMI-10.

### *In vitro* capture and phagocytosis assays

For flow cytometry-based capture assay, target cells were labelled with 5 µM of carboxyfluorescein succinimidyl ester (CFSE) from a CFSE cell proliferation kit (Invitrogen, C34554) for 5 min at room temperature, then washed three times with PBS containing 5 % FBS, and resuspended in RPMI with 5 % FBS before cells (4 x10^5^) were added to MDMs and cocultured at 37°C. After 2 h of co-culture, MDMs were extensively washed and analyzed by flow cytometry. Capture efficiency was determined as the percentage of CD11b^+^ cells containing CFSE-derived green fluorescence. For phagocytosis assay, target cells were labelled with 100 ng/ml of pHrodo Green STP ester (Invitrogen, P35369) pH 7.8 for 30 min at room temperature, resuspended in serum-free RPMI and then added to MDMs. After 2 h of co-culture at 37°C, MDMs were washed, collected and analyzed. Phagocytosis efficiency was determined as the percentage of CD11b^+^ cells containing pHrodo-derived green fluorescence.

### Coculture experiments of CD4^+^ T cells and autologous MDMs

CD4^+^ T cells were infected and after 2 days, washed, and maintained in culture for virus release during a 6 h incubation. Supernatants were separated from T cells by centrifugation (300 x *g*, 5 min), and fraction was added to MDMs for “cell-free” and “co-culture” infections, respectively. After the 6 h of incubation, MDMs were extensively washed to remove supernatants or T cells, and then cultured for 10 days. Media of MDMs were collected at specific interval of time for further analysis.

For the experiments involving Jasplakinolide (Jasp; Cayman Chemical, #102396-24-7), MDMs were pretreated with 5 µM of the inhibitor (or vehicle DMSO) for 1h. CD4^+^ T cells were subsequently cocultured with treated or untreated MDMs for 6 h in the presence or absence of Jasp. MDMs were washed extensively after the coculture and analyzed by flow cytometry for phagocytosis of pHrodo-labelled CD4^+^ target T cells by MDMs, as described above, or maintained in culture for 10 days for a replication kinetic study. Media of MDMs were collected at day 2 post coculture for measurement of infectious virus production by TZM-bl luciferase reporter assay (see below). To determine the effect of Jasp on virus production, infected CD4^+^ T cells were washed and incubated with Jasp or DMSO for 6 h, prior to cell supernatants being collected for quantification of virus production by HIV-1 p24 ELISA (XpressBio).

To assess the effect of reverse transcriptase inhibitor Zidovudine (AZT) or integrase inhibitor Raltegravir (Ral) on MDM infection, cells were pretreated with 5 µM AZT, 10 µM Ral or vehicle control DMSO for 2 h, and cocultured with infected CD4^+^ T cells in the presence of the drugs. After 6 h, MDMs were extensively washed to remove the T cells and drugs, and then cultured for another 2 days. Efficiency of MDMs infection was determined by flow cytometry analysis of intracellular Gag and production of infectious virus in MDMs-free culture supernatant using the TZM-bl assay.

### TZM-bl luciferase reporter assay

TZM-bl cells (2 x 10^4^ cells/well seeded in a 24-well plate the previous day) were inoculated with MDM-free culture supernatant for 6 h at 37°C, washed with PBS and maintained in DMEM-10. At 48 hpi, cells were lysed in cell culture lysis reagent (E153A, Promega) and analyzed for luciferase activity using a commercial kit (E1501, Promega).

### Confocal microscopy

SupT1 cells were infected with VSV-G pseudotyped GFP-expressing WT NL 4-3 ADA virus for 48 h and cocultured at a ratio of 4:1 with MDMs plated at 1,000 cells/well in an 8-well chamber slide (ibidi, #80806). After 2 h, MDMs were gently washed with PBS and fixed with 4% paraformaldehyde (PFA) for 30 min. Fixed MDMs were incubated for 2 h at 37°C in 5% milk-PBS containing anti-CD11b, a marker of macrophages. To detect p17, fixed MDMs were permeabilized in 0.2% Triton X-100 for 5 min, blocked in PBS containing 5% milk for 15 min, then incubated for 2 h at 37°C in 5% milk-PBS containing anti-p17 Abs which recognizes the mature matrix protein following Gag precursor processing by the viral protease but not the immature Gag precursor (73). Cells were washed and incubated with Alexa Fluor 594-coupled donkey anti-mouse IgG for 30 min at room temperature. Chamber slides were then washed with PBS and applied with DAPI solution (0.1 µg/mL in PBS) for 5 min, washed again and mounted using fluorescent mounting medium. Data were acquired using laser-scanning confocal microscope LSM-710.

### HEK 293T cell transfection

HEK 293T cells were transfected with appropriate plasmids using PEI. When applicable, the corresponding empty vectors were included in each transfection to ensure the same amount of transfected DNA in all conditions. For biochemical analyses involving the use of proteasomal and lysosomal inhibitors, MG132 (10 µM; Sigma-Aldrich, #474787) or Concanamycin A (ConA, 50 nM; Tocris Bioscience, #2656), respectively, were added to HEK 293T cells 36 h post-transfection. Cells were harvested for analysis 8 h thereafter.

### Western blotting

For SDS-PAGE and Western blotting analysis, cells were lysed in RIPA-DOC buffer (10 mM Tris pH7.2, 140 mM NaCl, 8 mM Na_2_HPO_4_, 2 mM NaH_2_PO_4_, 1% Nonidet-P40, 0.5 % sodium dodecyl sulfate, 1.2 mM deoxycholate) supplemented with protease inhibitors (cOmplete, Roche). Lysates were then mixed with equal volume of 2 x sample buffer (62.5 mM Tris-HCl, pH 6.8, 2 % SDS, 25 % glycerol, 0.01 % bromophenol blue, 5 % β-mercaptoethanol), and incubated at 37°C for 30 min as boiling was reported to cause aggregation of CD47 (82). Proteins from lysates were resolved on 15 % SDS-PAGE, transferred to nitrocellulose membranes, and reacted with primary antibodies. Endogenous CD47 was detected using a sheep polyclonal Ab, HA-tagged exogenous CD47 were detected using a rabbit mAb (clone C29F4). Membranes were then incubated with HRP-conjugated secondary Abs and proteins visualized by enhanced chemiluminescence (ECL).

### Co-immunoprecipitation assay

For co-immunoprecipitation studies of Vpu and CD47, transfected HEK 293T cells were lysed in CHAPS buffer (50 mM Tris, 5 mM EDTA, 100 mM NaCl, 0.5% CHAPS, pH7.2) supplemented with protease inhibitors. Lysates were first precleared by incubation with 40 µL of protein A-sepharose beads CL-4B (Sigma, #GE17-0963-03) for 1 h at 4°C and then incubated with mouse mAb anti-HA (clone 16B12) overnight. The following day, 40 µL of beads were added and samples were incubated for 2 h, washed five times with CHAPS buffer and analyzed by Western blotting.

### Flow cytometry

For analysis of CD47 surface expression on T cells, cells were washed with ice-cold PBS/EDTA (5 mM), stained at 4°C with anti-human CD47 or mouse IgG isotype control diluted in PBS/FBS (1%) for 30 min. Cells were then washed twice with PBS/FBS (1%) and fixed with 1% PFA. Apoptosis of target cells was evaluated using the Annexin V-PI detection kit (eBioscience, #88-8007-72) as per manufacturer’s protocol. For surface staining of MDMs, cells were washed with ice-cold PBS/EDTA (5 mM), detached with Accutase (Sigma-Aldrich, A6964), blocked in PBS/BSA (1%) /human IgG (blocking buffer) at 4°C for 20 min, and stained for 30 min with anti-human CD11b before additional washing and fixation with 1% PFA. For intracellular Gag staining, CD4^+^ T cells or MDMs were fixed and permeabilized using the Cytofix/Cytoperm kit (BD Biosciences) according to manufacturer’s instructions and stained with anti-Gag (KC57) at room temperature for 15 min, washed and resuspended in PBS/FBS (1%).

Flow cytometry data were collected on a Fortessa flow cytometer (BD Bioscience) unless specially specified. Cell sorting was conducted on a FacsAria (BD Bioscience). Analyses were performed using the FlowJo software, version 10.1 for Mac, BD Biosciences.

## ACKNOWLEDGMENTS

We thank members of our laboratory, especially Drs. Robert Lodge, Mariana G Bego and Sabelo Lukhele for helpful discussions and critical review of the manuscript; Frédéric Dallaire and Mélanie Laporte for technical support. We also thank Eric Massicotte and Julie Lord (IRCM Flow Cytometry Core) for assistance with flow cytometry; Odile Neyret and Myriam Rondeau (IRCM Molecular Biology Core) for their support with DNA sequencing; Martine Gauthier (IRCM clinic) for coordinating access to blood donors and all volunteers for providing blood samples. The following reagents were obtained through the NIH AIDS Reagent Program: SupT1 from D. Ablashi, TZM-bl from JC. Kappes and X. Wu, WITO (#11919), Zidovudine (#3485), Raltegravir (#11680) from Merck & Company, Inc. This study was supported by grants from the Canadian Institutes of Health Research (CIHR) (FDN 154324), the CIHR supported Canadian HIV Cure Enterprise 2 (CanCURE 2.0) (HB2 164064), and the Fonds de Recherche du Québec-Santé AIDS and Infectious Disease Network to É.A.C. L.C. was supported by studentships from the IRCM and Université de Montréal. É.A.C. is the recipient of the IRCM-Université de Montréal Chair of Excellence in HIV Research.

## SUPPLEMENTARY FIGURE LEGENDS

**Figure S1. Vpu downregulates overall levels of surface CD47.** Jurkat E6.1 cells were infected with VSV-G pseudotyped GFP-expressing NL 4-3 ADA (WT or dU) viruses and stained after 48 h with the indicated anti-CD47 mAbs (CC2C6 or B6H12) prior to flow cytometry analysis. **(Left)** Representative flow cytometry dot-plot graphs with indication of MFI values for infected (GFP-positive) and bystander cells (GFP-negative). **(Right)** Summary graphs of relative surface CD47 expression levels at 48 hpi with the indicated viruses (n=4). The percent MFI values were calculated relative to the respective GFP-negative cells. Statistical analysis was performed using Mann-Whitney U-test (*, P < 0.05), error bars represent SD.

**Figure S2. *In vitro* capture and phagocytosis assay controls. (A-D)** Characterization of target Jurkat E6.1 cell lines. **(A)** Flow cytometry histogram to validate CD47 surface expression levels in the indicated Jurkat cell lines; the *y* axis shows relative cell count for each population (normalized to mode) while the *x* axis shows fluorescence intensity of CD47. **(B)** Western blotting to assess total CD47 protein levels in the tested target Jurkat cell lines. **(C, D)** Summary graphs depicting the capture or phagocytosis of target Jurkat cell lines by MDMs as determined by CFSE- or pHrodo-labelling, respectively. Knockout of CD47 resulted in a better capture and phagocytosis of target cells by MDMs; n= 4 donors. **(E)** Representative flow cytometry dot-plots of Annexin V-propidium iodide (PI) staining of the indicated tested mock-infected or HIV-infected target cells for capture and phagocytosis assays (left); summary graphs for percentage of Annexin V^+^ population of tested target cells (right), n=5. **(C-E)** Statistical analysis was performed by Mann-Whitney U-test, (**, P < 0.01; *, P < 0.05; ns, nonsignificant, P > 0.05), error bars represent SD.

**Figure S3. Experimental controls. (A)** Infection of MDMs by WT NL 4-3 ADA (M-tropic) at MOI of 2, or WITO (T/F) at a MOI of 5; shown are the frequency of Gag^+^ MDMs from 3 donors at the indicated time points. **(B)** Infection levels of primary CD4^+^ T cells used for coculture with autologous MDMs; intracellular Gag staining was performed at 48 hpi and measured by flow cytometry. Statistical analysis was performed using Mann-Whitney U-test (ns, nonsignificant, P>0.05).

**Figure S4. Inhibition of reverse transcription and integration hinders the productive infection of MDMs**. MDMs were pretreated with raltegravir (Ral) or AZT, and cocultured with WT WITO-infected autologous primary CD4^+^ T cells in the presence of indicated drugs or vehicle (DMSO) for 6 h. After washing-off T cells and drugs, MDMs were cultured for 2 days and assayed for intracellular Gag by flow cytometry. Media of MDMs were collected at day 2 to infect TZM-bl cells as well. **(A)** Representative flow cytometry dot-plots showing the percentage of Gag^+^ cells at day 2 post coculture (top) and summary graph for MDMs from 3 distinct donors (bottom). **(B)** TZM-bl cells infected with media of MDMs collected at day 2 were assayed for Luc activity; shown are RLU detected with media collected from MDMs from 2 distinct donors; error bars represent SD.

**Figure S5. Generation and characterization of chimeric CD47.** Chimeric CD47 was generated by replacing the five membrane-spanning domains (MSDs) and cytoplasmic tail (CT) of human CD47 with the corresponding regions of mouse CD47. HEK 293T cells were co-transfected with plasmids encoding HA-tagged human, chimeric or mouse CD47 (HA-tagged at C-terminal), along with Vpu-expressing plasmid (pVpu). Whole cell lysates were analyzed by Western blotting. ECD: extracellular domain.

**Figure S6. Apoptosis control for phagocytosis assay. (Left)** Representative flow cytometry dot-plots of Annexin V-PI staining of the indicated tested target cells used for phagocytosis assay; **(Right)** summary graphs for percentage of Annexin V^+^ population of target cells, n=4, analyzed by Mann-Whitney U-test, (ns, nonsignificant, P > 0.05), error bars represent SD.

**Table S1. Oligonucleotides used in this study.**

